# Cell enlargement drives aging-associated proteome remodeling and shortens replicative lifespan

**DOI:** 10.64898/2026.02.15.706013

**Authors:** Michael C. Lanz, Manuel Hotz, Rachel Kroll-Ling, Eileen Wang, Jabob Kim, Joshua E. Elias, Daniel E. Gottschling, Jan M. Skotheim

## Abstract

The molecular and cellular basis of aging and its associated functional decline remains poorly understood. Even free-living microorganisms age and, in yeast, replicative aging shares key hallmarks with human cellular senescence, including progressive cell enlargement. Recent work has shown that chemical and genetic manipulations that increase cell size promote the onset of senescence in both yeast and human cells, suggesting that cell enlargement can drive some of the physiological changes associated with aging. Here, we quantitatively determined how cell enlargement contributes to age-associated physiology in yeast by combining automated aging technologies with quantitative proteomics. We find that the majority of aging-associated proteome remodeling can be recapitulated by genetically enlarging young proliferating cells. These enlarged cells exhibit accelerated proteome aging and shortened replicative lifespans, while smaller cells are longer-lived. While cell enlargement is the predominant factor driving proteome remodeling during aging, we also identified a minority of aging-specific molecular markers whose expression influences lifespan. Together, our results demonstrate that cell enlargement is a major driver of aging-associated proteome remodeling and influences lifespan independently of established aging factors such as extrachromosomal rDNA circles.

## Introduction

Cells undergo a progressive functional decline during aging whose molecular basis is poorly understood. For proliferative cells, repeated replicative cycles gradually erode the capacity to grow and divide, leading to a state of permanent cell cycle arrest known as senescence (Sharpless and Sherr, 2015; Hernandez-Segura et al., 2018). Studying aging in animals is difficult because lifespans are long and variable, making experiments slow and expensive. These drawbacks have motivated the study of aging in simpler microorganisms, such as the budding yeast *S. cerevisiae*, which exhibits many of the cellular phenotypes of aging observed in animals (Janssens and Veenhoff, 2016). In *S. cerevisiae*, replicative aging can be tracked across many generations in a simple, genetically tractable single-cell system, allowing for fast, well-controlled experiments that cleanly link environmental or genetic changes to lifespan (He et al., 2018; Steinkraus et al., 2008). *S. cerevisiae* divides asymmetrically, allowing larger “mother” cells to be distinguished from the smaller “daughter” cells that emerge from buds (Lindstrom and Gottschling, 2009). In yeast, the aging process is associated with several hallmark phenotypes, including the engagement of the general stress response (Hendrickson et al., 2018), rDNA instability and the accumulation of extra-chromosomal rDNA circles (ERCs) (Sinclair and Guarente, 1997), mitochondrial dysfunction (Guarente, 2008; Jazwinski, 2005), proteostasis decline (Moreno et al., 2019), and the accumulation of chitinous bud scars (Barton, 1950). Moreover, high-throughput studies have reported widespread age-associated changes in mRNA expression (Hendrickson et al., 2018; Yiu et al., 2008) and protein composition (Janssens et al., 2015; Lanz et al., 2024; Leutert et al., 2023). Despite the identification of these phenotypes that correlate with the progression of aging, few causal relationships have been established and the driving forces behind most aging-associated physiological changes remain unclear.

An often-overlooked feature of old yeast mother cells is their exceptionally large size. Cell size is typically tightly controlled within a given cell type, and aberrant cell size is often associated with pathological states (Ginzberg et al., 2015; Xie et al., 2022; Davies et al., 2022; Manohar and Neurohr, 2024). Pioneering work in yeast has found that birth size is predictive of replicative lifespan, with larger and smaller cells having shorter and longer lifespans, respectively (Yang et al., 2011). Moreover, yeast that are enlarged using temperature-sensitive alleles to temporarily block cell division undergo fewer replicative cycles before senescence (Neurohr et al., 2019). Despite these links between cell size and senescence, it has remained unclear how deviations in cell size would generally impact cell physiology, since cell composition was thought to scale proportionally with cell volume. However, a series of recent studies have overturned this paradigm by directly measuring gene expression as a function of cell volume. In every eukaryotic and prokaryotic cell type measured thus far, increasing cell size has been found to drive widespread changes in gene expression (Neurohr et al., 2019; Lanz et al., 2022; Terhorst et al., 2023; Lanz et al., 2024; Mäkelä et al., 2024; Tan et al., 2024; You et al., 2025). These cell size-dependent changes in gene expression are the consequence of a declining DNA-to-cell volume ratio (also known as “genome dilution”), since cell composition does not change when an increase in cell size coincides with a proportional increase in ploidy (Lanz et al., 2022, 2024). Collectively, these observations led us to further explore how the large size of aged yeast contributes to their altered physiology.

Here, we sought to quantify the extent to which cell enlargement contributes to the physiology of aged yeast cells. We previously showed that the proteome remodeling associated with increasing cell size was similar to that found in older cells (Lanz et al., 2024). However, this analysis relied on a batch-culture method of mother enrichment that was not of sufficient quality to accurately delineate aging-specific and cell size-specific proteomic changes. We therefore revisited this question by combining automated yeast aging technologies with modern proteomics. We find that the majority of age-associated proteome changes can be precisely recapitulated by genetically enlarging young yeast cells. Regressing out the contribution of enlargement to aging-associated gene expression changes revealed aging-specific molecular signatures. Finally, we show that genetically enlarged yeast exhibit an accelerated proteome-aging phenotype and shortened replicative lifespans, while smaller cells had prolonged lifespans. Together, our findings demonstrate that cell enlargement significantly contributes to the physiology of aged cells and influences lifespan independently of established aging factors such as ERCs.

## Results

### Replicative aging drives widespread changes in protein composition

To determine how the replicative aging process influences gene expression, we measured how proteome composition changes throughout the aging process. To do this, we used the <u>M</u>inistat <u>A</u>ging <u>D</u>evice (MAD) to isolate aged yeast mother cells (Hendrickson et al., 2018) and then comprehensively measured their proteome composition using TMT proteomics. The MAD captures magnetically-labeled yeast within a continuous culture vessel (**Figure 1A**). While retained within the vessel, the captured cells continue to proliferate, but their progeny are flushed from the device because they are not magnetically labeled (**Figure 1B**). To measure the progression of aging over time, we cultured captured mother cells for either 0, 24, or 48 hours, which corresponds to young, middle-aged, and old yeast, respectively (**Figure 1C**). The yeast proteomes from three independent aging time courses were then multiplexed and analyzed together using TMT proteomics.

**Figure 1:**
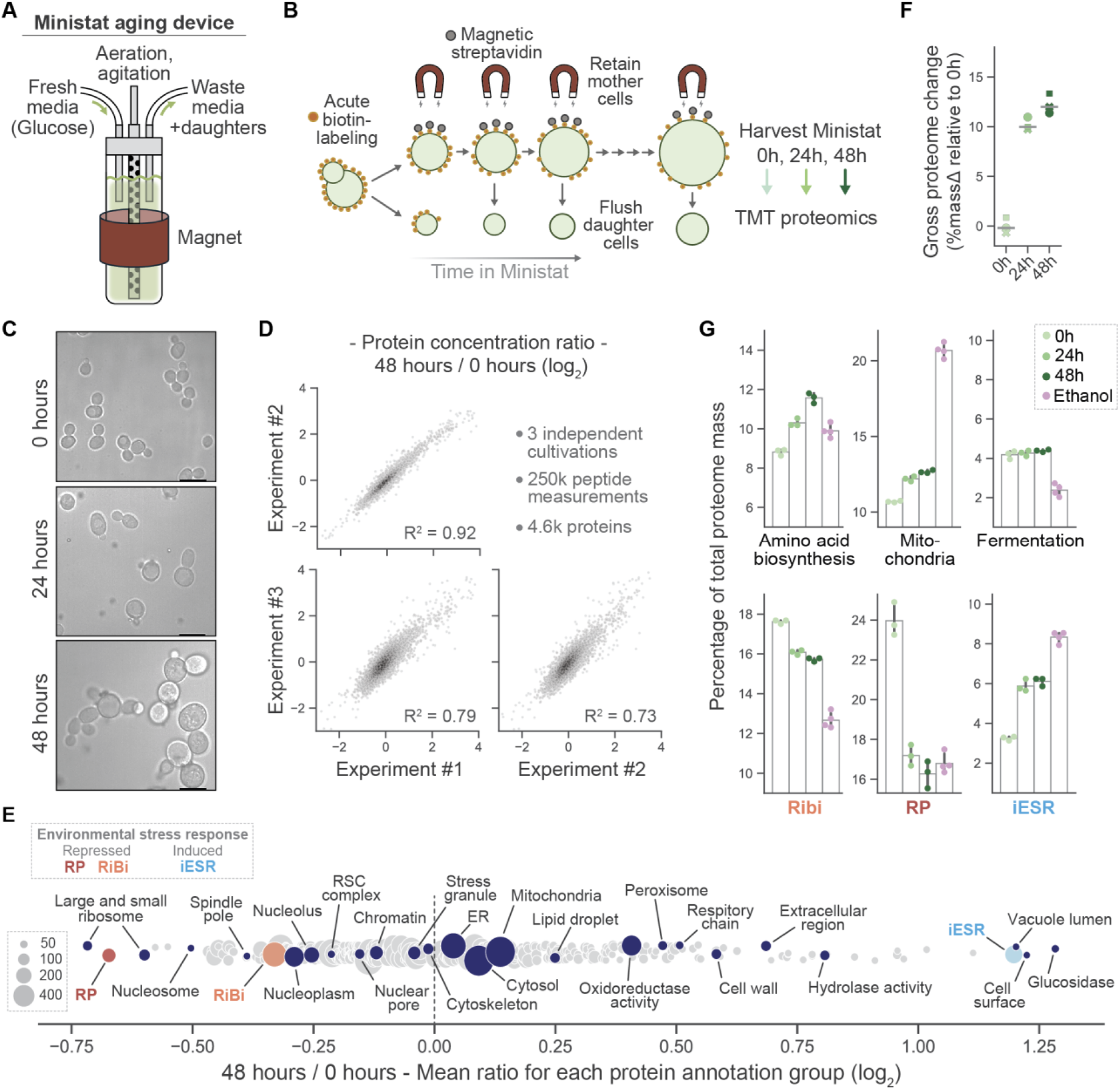
Comprehensive proteomic analysis of replicative aging using the Ministat Aging Device. A) Ministat Aging Device (MAD) is a continuous culture vessel that enables the separation of magnetic mother cells from non-magnetic daughter cells while maintaining a controlled and constant growth environment. B) Pipeline to measure the proteomes of replicatively aged yeast. Cultures are acutely labeled with biotin and magnetic streptavidin and then loaded into the MAD. Labeled cells are retained in the MAD via a magnet. Captured mother cells continue to divide, and the continuous flow of media through the vessel carries away unlabeled daughter cells. The extent of aging is controlled by the time captured cells are retained within the device. After three independent aging time courses, experiments were analyzed together using multiplexed proteomics. C) Images of yeast that were aged for different periods of time using the MAD. The 0 hour conditions represent yeast in log phase growth. Small dark dots in the 24 and 48 hour conditions are released (or free) magnetic beads. D) Correlation of age-associated proteome remodeling measured from three independent aging time courses. Each dot represents an individual protein, and its position reflects its relative change in concentration (log_2_) between yeast that were aged either 0 or 48 hours. Data density is shaded. E) Gene ontology analysis of aging-associated proteome changes. Each dot represents a GO “molecular function” or “cellular component” annotation. The size of each dot reflects the number of proteins with that GO annotation. The position of the dot along the x-axis indicates the mean protein concentration ratio (depicted in (D)) of all the proteins in that group. The y-axis is arbitrary jitter. ESR annotations were adopted from a previous study (Gasch et al., 2017). Ribosomal protein (RP), Ribosomal Biogenesis (RiBi), and Induced ESR (iESR) clusters are depicted. See Table S2 for all annotation groups. F) Approximation of total proteome mass change relative to the 0h condition for each cohort of aged yeast (see methods). The total variance between replicate 0h experiments was normalized to 0. Each point represents a measured proteome. Symbols correspond to the experiment number. G) Aging-associated changes in proteome mass allocation for the indicated protein groups. Each dot is a replicate proteome and color indicates the aging time in the MAD. “Ethanol” condition shows the allocation change in an experiment where yeast are grown at steady state using a nonfermentable carbon source (Lanz et al., 2024).

We found that protein composition changes substantially and reproducibly as a function of age (**Figure 1D and S1A; Table S1**). The relative concentrations of nearly a quarter of measured proteins (1,007 out of 4,446 proteins) increase or decrease by more than 50% as cells age. These changes did not correlate with the basal expression level (**Figure S1B**). Gene ontology analysis revealed that the concentrations of many nuclear and nucleolar proteins decrease with age, while the concentrations of vacuolar, respiratory, peroxisomal, and other proteins increase with age (**Figure 1E, Table S2**). We found that the most prominent gene expression signature associated with age was the environmental stress response (ESR) (**Figure 1E**), consistent with a previous transcriptomic study using a similar mother enrichment method (Hendrickson et al., 2018). In all, the aging process resulted in a gross proteome change of ~12% of relative protein mass (**Figure 1F**). Most of the proteome remodeling (by mass) corresponds to a shift from protein biosynthesis machinery in young cells, to more stress defense, amino acid anabolic, and mitochondrial proteins in older cells (**Figure 1G**). While the proteome remodeling associated with aging suggested an upregulation of respiratory metabolism, it was far less pronounced than what is observed after metabolic adaptation to a non-fermentable carbon source (**Figure 1G**). Importantly, while our measurements poorly correlated with a prominent published proteomic survey of yeast aging (Janssens et al., 2015), they strongly correlated with the age-associated proteome changes reported in a more recent study (Leutert et al., 2023) (**Figure S1C**). These newer datasets, which utilize ministats to maintain constant growth conditions during the aging process, establish a consensus proteomic signature of replicative aging in yeast.

### Cell enlargement during aging explains the majority of age-associated proteome changes

After finding that old and young yeast are compositionally different, we next sought to determine what aspect of the aging process drove these changes in gene expression. One prominent feature of aged yeast is their large size. With each round of division, the mother cell grows slightly larger (Yang et al., 2011). Indeed, we found that old cells (48h) were three times as large as young cells (**Figure 2A**). Because our recent work established that increasing cell size can remodel the proteome (Lanz et al., 2024, 2022), we hypothesized that the large size of old yeast could explain some age-associated changes in proteome composition.

**Figure 2:**
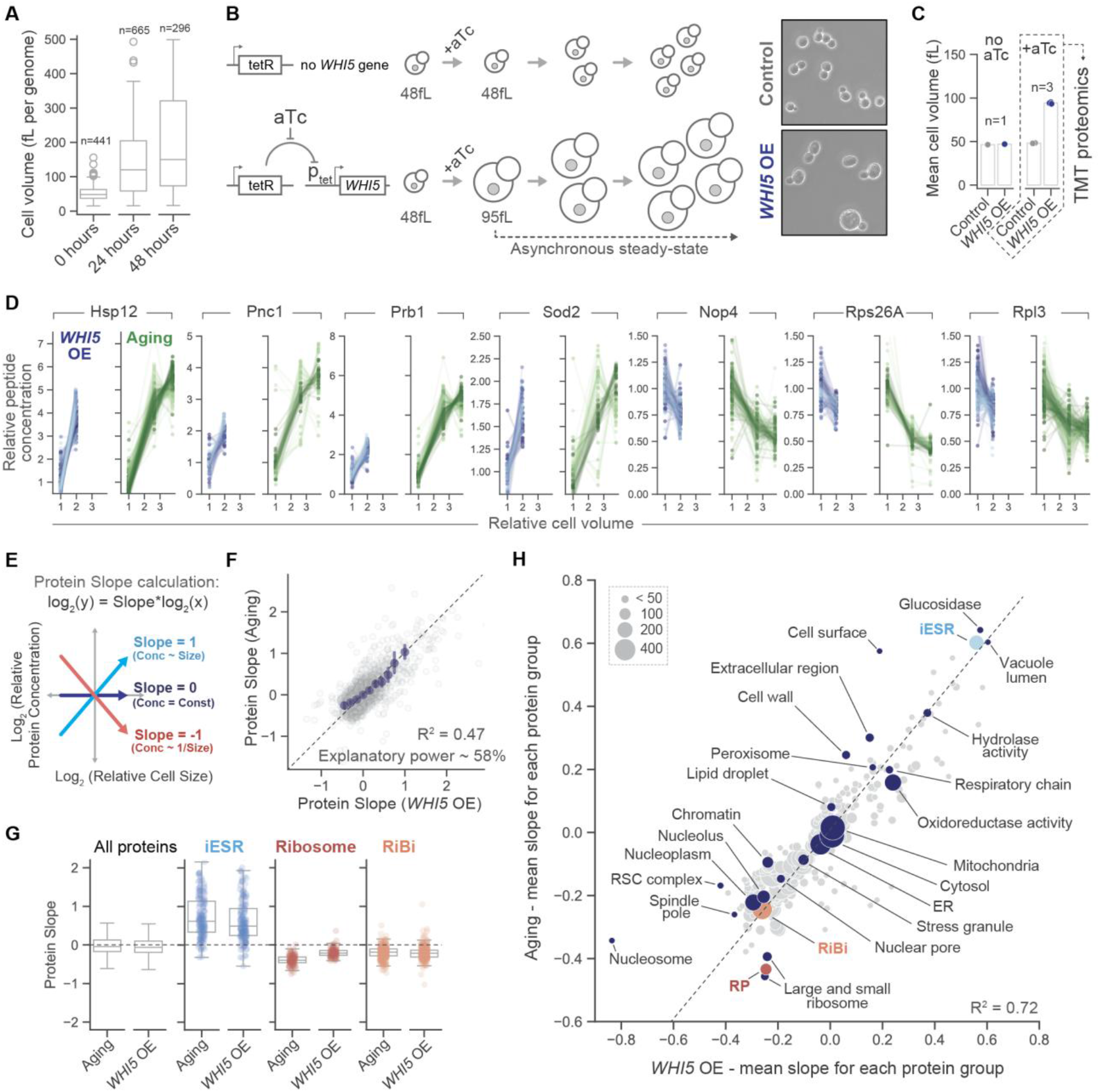
Cell enlargement explains the majority of aging-associated changes in proteome composition. A) Boxplots depicting the progressive enlargement of aged yeast cells. The sizes of individual cells were measured using images acquired immediately after removal from the MAD. Volumes were estimated using phase-contrast microscopy (see methods). B) An inducible system to generate large, proliferating yeast cells. A tetracycline-responsive promoter system was used to control the overexpression of the G1/S inhibitor gene, *WHI5*. Overexpression of *WHI5* results in large cells that are still dividing. Proteomes were sampled ~12 generations after induction, by which point the cultures had reached a new steady state cell size and doubling time (see Figure S2). *C) WHI5* overexpression roughly doubles the set-point size around which cells proliferate asynchronously. Mean cell size was determined using a Coulter counter. D) Relative peptide concentration changes for the indicated proteins measured across the indicated conditions by multiplexed TMT mass spectrometry. Each dotted line represents an independent peptide measurement corresponding to the indicated protein. Shades of blue and green show peptides measured from each biological replicate experiment. All concentration changes are measured relative to “Control” or “0h”. Relative cell volume was determined using the mean of the data depicted in (A) and (C). E) Derivation of the protein slope value. Protein slopes describe the size-scaling behavior of each individual protein. Proteins with a slope of 0 maintain a constant cellular concentration regardless of cell volume. A slope value of 1 corresponds to an increase in concentration that is proportional to the increase in volume and of −1 to dilution so that the concentration is inversely proportional to volume. F) Correlation of protein slopes derived from genetically enlarged and replicatively aged cells. The identity line is dashed. Blue dots are x-binned data and error bars represent the 99% CI. “Explanatory power” is an R^2^ value that is corrected by the measured variance between biological replicate experiments in Figure 1D. G) Box plots showing how the yeast stress response is expressed proportionally to cell size in both genetically enlarged and replicatively aged cells. Each individual dot is the scaling slope for an individual protein of the indicated category. ESR annotations were adopted from a previous study (Gasch et al., 2017). H) Gene ontology analysis comparing the proteome changes associated with aging and *WHI5* overexpression. Each dot represents a GO “molecular function” or “cellular component” annotation. The size of each dot reflects the number of proteins with that GO annotation. The position of the dot indicates the mean protein slope value of all the proteins in that group. See Table S2 for all annotation groups.

To measure how cell enlargement influences gene expression, we used a genetic system to enlarge proliferating populations. Unlike our previous methods to manipulate cell size (Lanz et al., 2024), this system generates proliferating cells of a size comparable to aged mother cells. In brief, we expressed the G1/S regulatory protein Whi5 from a synthetic TET promoter that is conditionally activated by anhydrotetracycline (aTc) (Azizoglu et al., 2021) (**Figure 2B and S2**). Overexpression of Whi5 (“*WHI5* OE”) doubled the mean cell size of the population (**Figure 2C and S2A**) while only slightly reducing the exponential population doubling time (**Figure S2B**). Importantly, this enlarged population is still dominated by newly budded daughters and young mother cells. Older cells are diluted out by exponential growth and are therefore exceedingly rare in the population. Proteomic analysis of genetically enlarged yeast revealed widespread changes in gene expression that strongly correlated with our previous measurements of size-dependent proteome remodeling in G1-arrested yeast and single deletion cell size mutants (**Figure S3**). Thus, similar to the comparison of old and young yeast, enlarged proliferating yeast are also compositionally different from their normally-sized counterparts.

We next compared size- and age-associated changes in proteome composition. If a given protein’s concentration were to increase as a function of cell size, we would expect it to also progressively increase as yeast age. Indeed, when comparing the loading-normalized peptide ion intensities, which are a direct proxy for relative peptide concentration change across conditions, we find the protein concentrations for several age-associated proteins (Anderson et al., 2003; Fabrizio et al., 2003) changed similarly as a function of cell volume in both genetically enlarged and aged cells (**Figure 2D**). To more quantitatively compare size- and age-associated proteome remodeling, we calculated how the concentration of each protein changes as a function of cell size using a simple regression (**Figure 2E**; Protein “Slope”), as described previously <u>(Lanz et al.,</u> 2024, 2022). Proteins with a slope of 0 maintain a constant cellular concentration regardless of cell volume. A slope value of 1 corresponds to an increase in concentration that is proportional to the increase in volume while −1 represents dilution so that the concentration is inversely proportional to the volume. When the aging- and genetic enlargement-derived protein slopes are directly compared, the resulting correlation is strong (Pearson *r* = 0.73) and lies on the identity line (R^2^ = 0.47) (**Figure 2F; Table S1**). Other methods of cell enlargement produced similarly strong correlations (**Figure S3**), which indicates that the proteome changes are truly size-driven rather than manipulation-specific. The most prominent gene expression signature associated with aging, the ESR, is induced in close proportion to cell size in both aged and genetically enlarged yeast (**Figure 2G**). Grouping proteins by GO annotation yielded an even stronger correlation (R^2^ = 0.72) (**Figure 2H**). If we use the measurement noise between our biological replicate experiments as a benchmark for the maximum achievable correlation (an average R^2^ of 0.81 for our 3 replicate experiments in Figure 1D), then the correlation between size- and age-dependent proteome changes (R^2^ of 0.47; **Figure 2F**) indicates that cell enlargement recapitulates most of the compositional changes observed in aged yeast.

### Identification of aging-specific compositional changes

Since cell enlargement can account for most of the gene expression changes associated with replicative aging, then regressing out the “cell size effect” from our aging dataset should reveal aging-specific compositional changes. To do this, we subtracted our aging-derived slope values by the *WHI5* OE-derived slopes, which reduced the overall variance in the dataset (**Figure 2F & 3A**). Gene ontology comparison of uncorrected and size-subtracted aging protein slopes revealed a subset of composition sectors with aging-specific changes (**Figure 3B**), most notably cell surface, vacuolar, ribosomal, and histone proteins (**Figure 3C**). Interestingly, while we observe a “loss” of histone protein in our aged cells (Hu et al., 2014), the decrease in histone concentration was relatively less than in genetically enlarged cells (**Figure 3C**), where changes in histone concentration closely match the change in genome concentration (Claude et al., 2021). This regression analysis demonstrates the importance of accounting for cell enlargement in order to identify genuine molecular markers of aging.

**Figure 3:**
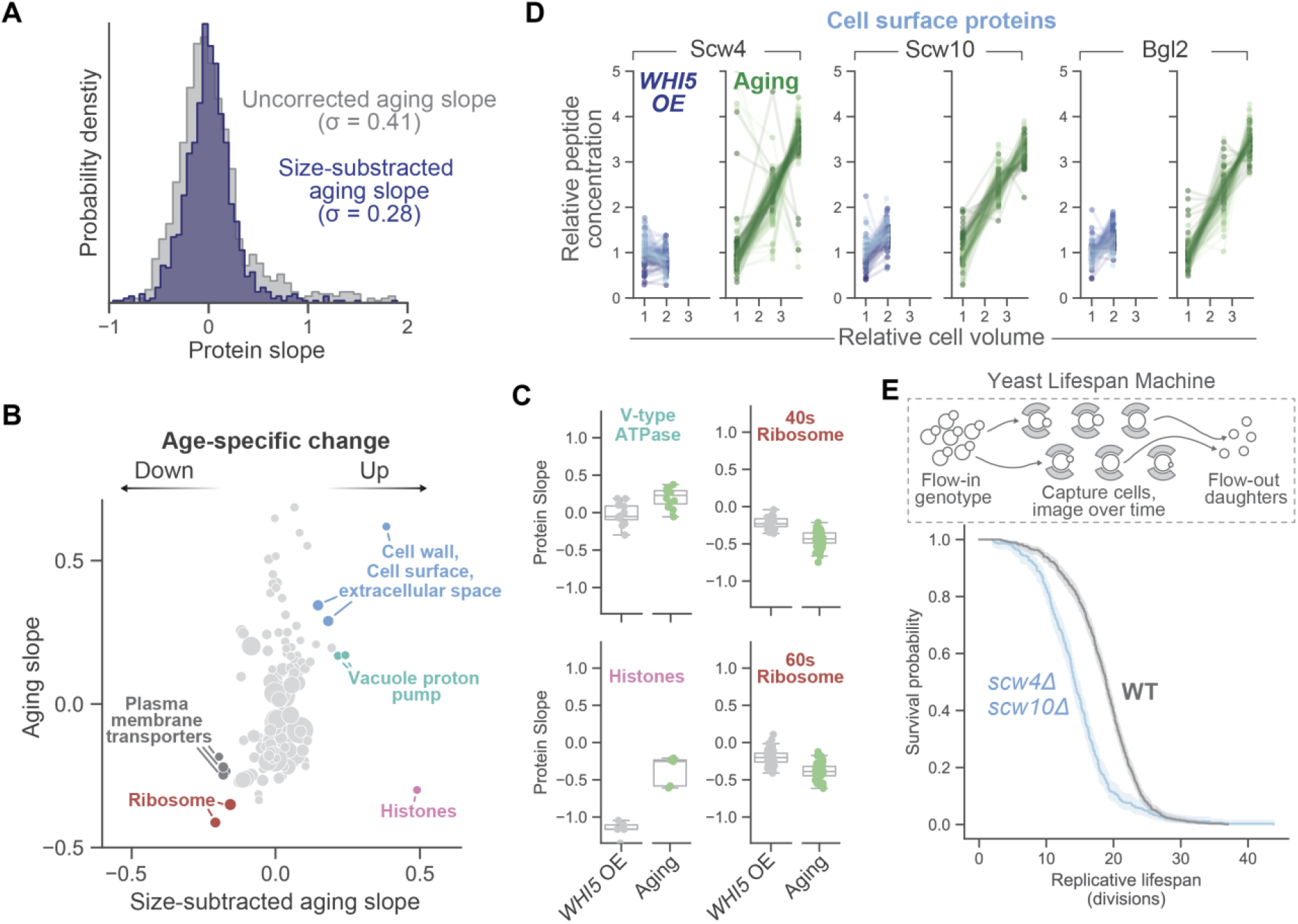
Regressing out the cell size effect reveals aging-specific compositional changes. A) Distributions of age-associated protein slope values describing the concentration change for individual proteins before and after subtracting the effect of cell size. σ denotes the standard deviation of the distribution. Size-subtracted aging slope values were calculated by subtracting the “*WHI5* OE” slope from the “Aging” slope (Figure 2F; Table S3). B) Gene ontology analysis comparing aging-associated and aging-specific proteome changes. Each dot represents a GO “molecular function” or “cellular component” annotation. The size of each dot reflects the number of proteins with that GO annotation. The position of the dot indicates the mean protein slope value of all the proteins in that group. C) Box plots depicting the average scaling behavior of the indicated protein groups. Each individual dot is the scaling slope for an individual protein. D) Relative peptide concentration changes measured across the indicated conditions by multiplexed TMT mass spectrometry. Each dotted line represents an independent peptide measurement corresponding to the indicated protein. Shades of blue and green show peptides measured from each biological replicate experiment. All concentration changes are measured relative to “Control” and “0h”. E) Top panel: The Yeast Lifespan Machine is an automated method for measuring replicative lifespan of a desired genotype (Thayer et al., 2022). Bottom panel: Kaplan-Meier lifespan curves for the indicated genotypes. 95% confidence interval is shaded.

We next examined whether the aging-specific proteome changes we identified could be functionally important. The proteins with the strongest aging-specific upregulation were those associated with the cell surface (**Figure 3B**). To test whether the expression of these cell surface-related proteins was functionally important for aged yeast, we deleted two paralogous cell surface proteins whose expression was specifically upregulated in the old yeast: the putative glucanases Scw4 and Scw10 (**Figure 3D**). To measure how the lifespan of a *scw4Δscw10Δ* double mutant strain changed relative to WT, we utilized the Yeast Lifespan Machine (YLM), an automated imaging platform that can simultaneously determine the replicative lifespans of thousands of individual yeast cells (Thayer et al., 2022). In brief, the device captures individual cells on a flow cell and successively images them over a period of days to measure the total number of progeny produced from each captured cell and thus determine their replicative lifespan (**Figure 3E**). Using the YLM, we found that our *scw4Δscw10Δ* double mutant had a significantly shorter lifespan than our wild-type strain (**Figure 3E**), despite showing no observable fitness or cell size defect in batch culture. Thus, the expression of Scw4 and Scw10 is necessary to achieve a normal lifespan in budding yeast.

### Cell size accelerates aging independent of extrachromosomal rDNA circle accumulation

Because cell enlargement drives age-associated changes in gene expression (**Figure 2**), we next examined whether enlarging young yeast accelerates the aging process (**Figure 4A**). First, we used the MAD to monitor the progression of aging in enlarged yeast using proteome composition as a proxy for age. Similar to our Whi5 OE strain, a *cln3Δ* mutant, which is ~50% larger than WT, exhibited aging-associated proteome changes in log phase culture (**Figure S4**). Interestingly, after aging this mutant strain 24 and 48 hours, it possessed a proteome phenotype significantly older than its WT counterpart (**Figure S4**). Indeed, both *cln3Δ* and Whi5 OE strains exhibited dramatically reduced lifespans, consistent with their proteome’s resembling that of aged cells (**Figure S4**). Conversely, *whi5Δ* mutants, which are slightly smaller than WT, had extended lifespans (**Figure 4B and 4C**). This finding is consistent with previous work (Yang et al., 2011, 2018), though the lifespan lengthening effects of a *whi5Δ* mutant are contested (Crane et al., 2019). In support of our measurements of *whi5Δ*’s extended lifespan, the proteomes of the smaller *whi5Δ* mutants anticorrelate with the changes observed in the larger *cln3Δ* (**Figure S4**) (Lanz et al., 2024). The finding that small and large cell size mutants can increase and decrease replicative lifespan, respectively, supports a previous pioneering study linking cell size and replicative lifespan (Yang et al., 2011). Taken together, our data indicate that cell enlargement can accelerate the progression of replicative aging.

**Figure 4:**
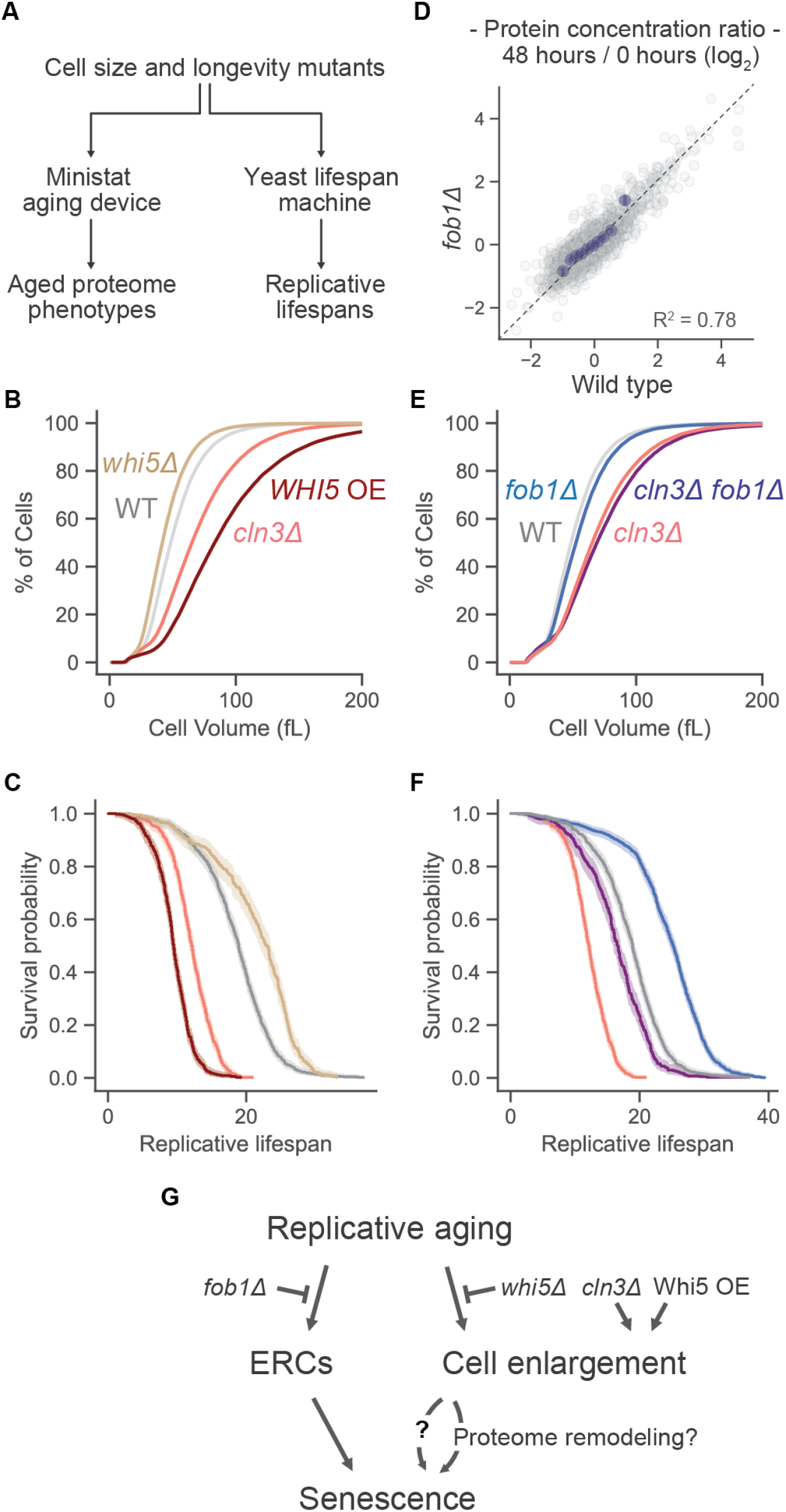
Cell size influences lifespan independently of ERCs. A) Schematic illustrating the plan to measure the progression of aging in cell size and longevity mutants using proteomics and imaging. B) Cumulative distributions of cell volume for the indicated genotypes. Cell volumes were determined from log-phase cultures using a Coulter counter. C) Kaplan-Meier lifespan curve for the genotypes highlighted in (B). Colors correspond to genotype labels in (B). 95% confidence interval is shaded. D) Correlation of aging-associated proteome changes in WT and *fob1Δ* mutant cells. Each dot represents an individual protein, and its position reflects its relative change in concentration (log_2_) between yeast that were aged either 0 or 48 hours. The identity line is dashed. Blue dots are x-binned data and error bars represent the 99% CI. E) Cumulative distributions of cell volume for the indicated genotypes. Cell volumes were determined from log-phase cultures using a Coulter counter. F) Kaplan-Meier lifespan curve for the genotypes highlighted in (E). Colors correspond to genotype labels in (E). 95% confidence interval is shaded. G) A model for how different mutations impact the accumulation of ERCs or cell enlargement and how these phenotypes independently contribute to replicative senescence in yeast.

After confirming the link between cell size and replicative aging, we next sought to determine how cell size interacts with another established determinant of yeast lifespan, the accumulation of extrachromosomal rDNA circles (ERCs). As yeast mothers perform successive rounds of DNA replication, circular copies of the highly repetitive rDNA locus can be ejected from the genome via homologous recombination (Sinclair and Guarente, 1997). The accumulation of ERCs is strongly associated with replicative senescence, and mutants that prevent (*e*.*g*., *fob1Δ*) or promote (*e*.*g*., *sir2Δ*) the accumulation of ERCs increase and decrease lifespan, respectively (Defossez et al., 1999; Kaeberlein et al., 1999). To determine whether ERC accumulation influenced the proteome changes associated with aging, we measured the progression of aging in the long-lived *fob1Δ* mutant using the MAD. We found that the proteomes of *fob1Δ* mutants are nearly indistinguishable from WT after 24 hours of aging (**Figure 4D and S4**), suggesting the presence or absence of ERCs does not influence how gene expression changes with age. To test how the lifespan-influencing effects of ERCs interacted genetically with cell enlargement, we generated a *fob1Δcln3Δ* double mutant. Interestingly, we found that the median lifespan of the *fob1Δcln3Δ* strain was similar to WT (**Figure 4E and 4F**). Together with the fact that *fob1Δ* does not greatly affect cell size or proteome composition, this non-epistatic phenotype indicates that the lifespan-influencing effects of cell enlargement and ERCs are mostly independent of one another (**Figure 4G**).

## Discussion

Here, we identify cell enlargement as a major driver of age-associated physiological change. We show that the majority of proteome remodeling observed during replicative aging can be sufficiently explained by increases in cell size throughout aging. This finding provides a unifying framework for understanding the driving force behind hallmark features of aging in yeast, like histone “loss” (Hu et al., 2014) and ESR activation (Hendrickson et al., 2018). Our results argue that replicative aging is not solely driven by stress or the accumulation of molecular damage but is instead strongly shaped by a gene expression program driven by cell enlargement.

Though the aging process coincides with widespread proteome remodeling, it remains unclear which changes in composition causally contribute to replicative senescence. Compositional changes that accompany cell enlargement, and therefore also accompany aging, could potentially relate to previously proposed causes of yeast mortality, such as mitochondrial dysfunction (Guarente, 2008) or a decline in proteostasis (Moreno et al., 2019). In support of this notion, we do report size-dependent changes in mitochondrial composition, as well as in the concentrations of over a dozen protein folding chaperone proteins (see Table S1). It is also still unclear how the most prominent gene expression program associated with cell enlargement, the ESR (**Figure 2G**), influences lifespan. On one hand, the ESR could be activated by some as-of-yet unknown stress related to enlargement. Alternatively, size-dependent activation of the ESR could represent an evolved mechanism to passively prepare smaller daughters for the stress that subsequently comes with aging. Regardless of its causality, a recent study demonstrated that ESR induction in large, arrested yeast cells suppresses a decline in their viability (Terhorst et al., 2023), which suggests that the ESR in large, aged cells could be beneficial.

By accounting for size-dependent effects, we were also able disentangle aging-specific molecular changes from those that passively arise as a consequence of cell enlargement for the first time. Although only a few sectors of cell composition were specifically responsive to age, each yields critical insight into the physiology of aged cells. For example, the age-specific increase in the vacuolar proton pump (**Figure 3C**) could represent a compensatory mechanism to counteract an aging-dependent decline in vacuole pH (Hughes and Gottschling, 2012). The dramatic upregulation of cell surface-related proteins (**Figure 3D**) might reflect a strategy to cope with the bud scars that accumulate with successive divisions. Aging also appears to disrupt the stoichiometric relationship between histones and the genome because aged cells have more histone proteins relative to the amount of DNA than young cells (**Figure 3C**). The increased number of histones relative to the genome would plausibly contribute to senescence given the toxicity of excess histones (Gunjan and Verreault, 2003). Finally, ribosome concentrations decrease significantly more during aging than what would be expected by cell enlargement alone (**Figure 3C**), which could reflect an aging-specific engagement of ribophagy (Kraft et al., 2008). We anticipate future work will determine how these aging-specific compositional changes arise, whether they ultimately influence lifespan, and how they relate to other biomarkers of aging that are more specific than general changes in protein or mRNA concentration (Hughes et al., 2016; Mouton et al., 2020; Gutierrez and Tyler, 2024; Mobaraki et al., 2026).

Large cell size could also influence replicative senescence in ways other than driving changes in proteome composition. Previous work has shown that excessive cell enlargement leads to cytoplasmic dilution and impairs transcriptional induction in yeast (Neurohr et al., 2019). This decline in transcriptional responsiveness could potentially prevent the induction of cyclin expression in large mother cells and thus drive permanent G1 arrest (Neurohr et al., 2018). How transcriptional induction is compromised in large cells is still unclear, though it may relate to the sublinear size-scaling of RNA polymerase II activity (Swaffer et al., 2023). Future work will test how these and other size-dependent physical constraints on cell physiology limit replicative lifespan in aging cells.

Although replicative aging in yeast and cellular senescence in mammals occur in distinct contexts, several striking parallels suggest that cell enlargement may represent a conserved driver of senescence induction. Much like replicative senescence in yeast, large size is a near-universal marker of mammalian senescence (Hernandez-Segura et al., 2018), and increasing cell size has been found to drive many senescence-associated proteome changes in cultured mammalian cells (Cheng et al., 2021; Lanz et al., 2022). Moreover, excessive cell enlargement caused by CDK4/6 inhibitors promotes durable cell cycle exit even in the absence of overt DNA damage, and interventions that prevent the enlargement of CDK4/6i-treated cells preserve their proliferative capacity (Neurohr et al., 2019; Lanz et al., 2022; Crozier et al., 2023; Foy et al., 2023; Manohar et al., 2023; Wilson et al., 2023). These observations, together with the work in yeast presented here, position cell enlargement as a conserved driver of eukaryotic senescence.

## Methods

### Yeast strain construction and cell culture

Yeast strains were constructed using standard transformation protocols and antibiotic selection markers. All yeast in this study were cultured using synthetic complete media supplemented with 2% glucose (SCD) unless otherwise indicated. To induce transcription from the *P*_*Tet*_*-WHI5*, 50 ng/mL of anhydrotetracycline (aTc) was added to each culture flask to relieve TetR repression. In the case where *WHI5* OE yeast were measured using the lifespan device, the media in the flow cell was supplemented with 50ng/mL aTc to maintain *WHI5* OE.

### Ministat Aging Device (MAD)

Cells were aged in the MAD system as previously described (Hendrickson et al., 2018; Hotz et al., 2022). Briefly, cells were grown in SCD medium with three modifications compared to regular SCD: YNB was replaced with biotin-free YNB (Sunrise Science), 40 nM 7,8-diaminopelargonic acid (DAPA) was added to compensate for the lack of biotin, and 2% methyl-alpha-D-mannopyranoside (Sigma-Aldrich) was added to prevent cell aggregation during the experiment. Cells were grown for 24 hours at OD < 0.05 and 5 ODs were harvested for each experiment. Cells were washed with PBS and incubated with 2.5 mg Sulfo-NHS-LC-LC-Biotin (Sigma-Aldrich) for 30 min at room temperature. Biotinylated cells were then washed three times with medium and added to flasks with medium for 4-5 hours. Cells were then incubated with 20 µL of Dynabeads MyOne Streptavidin C1 magnetic beads (Thermo Fisher) for 10 min at room temperature. Cells were then washed on a DynaMag-2 magnet (Invitrogen) and loaded into a MAD pre-filled with medium. After binding to the magnet for 5 minutes, media pumps were turned on. Cells were either harvested after 24 or 48 hours as indicated. For timepoints later than 24 hours, MADs were washed after 24 hours to avoid overcrowding.

### Replicative lifespan measurements

Replicative lifespan was measured in the Yeast Lifespan Machines (YLM) as described in (Hotz et al., 2022; Thayer et al., 2022). Briefly, cells were grown for 24 h at OD below 0.05 in SCD medium and then loaded into microfluidics devices made from polydimethylsiloxane (PDMS) with catcher structures designed to retain aging mother cells and continuously washing away daughter cells. These devices had 24 parallel channels and each channel contained ~1,000 “DetecDiv”-style catchers (Aspert et al., 2022). Brightfield images were collected every 15 min for 96 h and analyzed using an automated computer vision pipeline, capable of counting the number of divisions and the terminal outcome of a cell’s lifespan (death vs. censoring/washout). Median lifespan was determined using the Kaplan-Meier estimate of the survival function.

### Cell volume measurements

Cell volumes were measured using either a Z2 Coulter counter (Beckman) or microscopy. We used microscopy to measure relative changes in cell volume of aged cells because Calico Labs did not have a functioning Coulter counter. We have previously shown that relative changes in volume measured by microscopy closely correspond to those measured via Coulter counter (Lanz et al., 2024). For cell volume measurements estimated from imaging, we utilized an existing algorithm in Cell-ACDC (Padovani et al., 2022). Cell-ACDC was used to segment the cell boundaries with the convolutional neural network algorithm YeaZ (Dietler et al., 2020). The volumes of mother and buds were counted separately. Small buds were excluded from the analysis (early S phase) to better estimate the relative changes in cell volume per genome.

### Sample preparation for mass spectrometry analysis

Yeast cultures were pelleted by centrifugation or collected on a 0.2-µm disposable filter. Pellets were resuspended in 300 µl of yeast lysis buffer (50 mM Tris, 150 mM NaCl, 5 mM EDTA and 0.2% Tergitol, pH 7.5 + a cOmplete ULTRA Tablet) with 700 µl of glass beads. Lysis was performed at 4 °C in a Millipore Fastprep24 (4 cycles with the following settings: 6.0 m s−1, 4× 40 s). Cell lysates were cleared by centrifugation at 12,000g for 5 min at 4 °C. Protein concentration was quantified using a Pierce BCA Protein Assay Kit (product no. 23255). Lysates were then denatured/reduced in 1% sodium dodecylsulfate (SDS) and 10 mM dithiothreitol (15 min at 65 °C), alkylated with 5 mM iodoacetamide (15 min at room temperature) and then precipitated with three volumes of a solution containing 50% acetone and 50% ethanol (on ice for 10 min). Proteins were re-solubilized in 2 M urea, 50 mM Tris-HCl and 150 mM NaCl, and then digested with TPCK-treated trypsin (50:1) overnight at 37 °C. Trifluoroacetic acid (TFA) and formic acid were added to the digested peptides for a final concentration of 0.2% (pH ~2). Peptides were desalted with 50-mg Sep-Pak C18 columns (Waters).

TMT-labeled peptides were fractionated using either an offline HPLC or the Pierce High pH Reversed-Phase Peptide Fractionation kit. HPLC fractionation was performed on the primary aging and enlargement dataset to achieve maximal depth (24 fractions). For cell size and longevity mutant experiments, the fractionation kit was used. The eight default fractions were either injected separately or pooled back into four fractions (1-5, 2-6, 3-7 and 4-8). In all cases, dried peptides were reconstituted in 0.1% TFA. Peptide concentrations were determined using a Nanodrop before injection. See Table S5 for a complete summary of all proteome experiments.

### LC-MS/MS data acquisition

TMT-labeled peptides were processed as described previously. Briefly, they were resuspended in 0.1% formic acid and analyzed on a Fusion Lumos mass spectrometer (Thermo Fisher Scientific, San Jose, CA) equipped with a Thermo EASY-nLC 1200 LC system (Thermo Fisher Scientific, San Jose, CA). Peptides were separated by capillary reverse phase chromatography on a 25 cm column (75 µm inner diameter, packed with 1.6 µm C18 resin, AUR2-25075C18A, Ionopticks, Victoria Australia) and introduced with 180-min stepped linear gradient at a flow rate of 300 nL/min. The gradient steps were: 6–33% buffer B (0.1% (v:v) formic acid in 80% acetonitrile) for 145 min, 33-45% buffer B for 15 min, 45–95% buffer B for 5 min and maintained at 90% buffer B for 5 min. The column temperature was heated to 50 °C throughout the procedure. Xcalibur software was used for the data acquisition. Survey scans were acquired in the Orbitrap (centroid mode) over the range 380–1,400 m/z with a resolution of 120,000 (at m/z 200). For MS1, the normalized AGC target (%) was 250 and the maximum injection time was 100 ms. Ions were fragmented by collision-induced dissociation (CID) with normalized collision energies of 34. The isolation window was set to a 0.7-m/z window. MS2 was acquired in the ion trap mass analyzer with the scan rate set to ‘Rapid’, the normalized AGC target was set to ‘Standard’, and maximum injection time to 35 ms. Dynamic exclusion of the sequenced peptides was set to 30 s. The maximum duty cycle time was set to 3 s. Relative changes in peptide concentration were determined at the MS3 level by isolating and fragmenting the five most dominant MS2 ion peaks.

### Spectral searches

All raw files were searched using the Andromeda engine embedded in MaxQuant (v2). Reporter ion MS3 search was conducted using TMT10-plex or TMT16-plex settings depending on the experiment. Variable modifications were oxidation (M) and protein N-terminal acetylation. Carbamidomethyl (C) was a fixed modification. The number of modifications per peptide was capped at five. Digestion was tryptic (proline-blocked). Database search used the UniProt “_YEAST” proteome (Saccharomyces cerevisiae). The minimum peptide length was 7 amino acids. 1% FDR was determined using a reverse decoy proteome. See Table S5 for a complete summary of all proteome experiments.

### Mass spectrometry data analysis

Mass spectrometry analysis was performed as described previously (Lanz et al., 2022). Briefly, protein-level measurements were assembled from MaxQuant’s evidence.txt output file. Each row of the evidence.txt file represents an independent peptide and its corresponding MS3 reporter ion measurements. Peptides were first filtered for decoy and contaminants. Peptides without a signal in any of the TMT channels were excluded. For each individual peptide measurement (*i*.*e*., each row in the evidence table), the fraction of ion intensity in each TMT channel was calculated by dividing the “Reporter ion intensity” column by the sum of all reporter ion intensities for that peptide. To correct for loading differences between the TMT channels, each reporter ion channel was then normalized by dividing the fraction of ion intensity in each channel by the median fraction for all measured peptides (*i*.*e*., the median value for each column). This normalization scheme ensures that each individual peptide measurement is equally weighted when correcting for loading error. This loading-corrected value was used as a measure of relative peptide concentration across all TMT channels. See Table S5 for a complete summary of all proteome experiments.

Protein slope calculation was performed as described previously (Lanz et al., 2022). Briefly, individual peptide measurements were consolidated into a protein-level concentration measurement using Python’s groupby.median. Peptides with the same amino acid sequence that were identified as different charge states or in different fractions were considered independent measurements. We summarized the size-scaling behavior of individual protein as a slope value derived from a log-log regression of its relative changes in concentration as a function of mean cell volume/. Each protein slope is based on the behavior of all of its detected peptides.

For a given protein, we calculated its slope as follows:

*y*_*i*_ = relative signal in the *i*th TMT channel (median of all peptides measured for this protein).

*x*_*i*_ = relative cell volume corresponding to the sample in the *i*th TMT channel.

The protein slope value was determined from a linear fit to the log_2_(transformed data) using the equation:

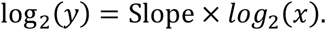

Variables were log_2_(transformed) so that a slope of 1 corresponds to an increase in protein concentration that is proportional to the increase in volume and a slope of −1 corresponds to 1/volume dilution.

### Regression analyses

Regression plots used throughout the manuscript were generated using seaborn’s “regplot” package in Python. Grayed dots represent all data points in the regression analysis. Each bin is represented by a dark-blue dot. Each dot represents a bin’s mean y-axis value and its error bar is the 99% confidence interval (CI) for the y coordinate. Pearson’s r and P values for correlation analyses were calculated using SciPy’s pearsonr module in Python. R^2^ values were calculated using “r2_score” from scikit-learn.

### Gene ontology analyses

1D annotation enrichment analysis was performed as described previously (Cox and Mann, 2012) using the Perseus platform. Each protein was annotated according to its “Molecular Function” and “Cellular Component” terms. 1D enrichment analysis was used to group proteins by shared annotation and determine the mean and median quantitation value for that group. For the GO analyses in Figure 2 and 3, 1D annotation enrichment analysis was performed for each axis and then plotted together.

### Proteome mass fraction calculation

To calculate the proteome mass fraction of each mapped protein, we utilized the peptide feature information in MaxQuant’s evidence.txt output file. To calculate proteome mass fractions, the MS1 precursor ion intensity of each peptide measured (the “Intensity” column in the evidence.txt table) was distributed between the individual MS3 reporter channels according to the loading-normalized value described above. Protein-level ion intensities were then calculated for each TMT channel by summing together all peptide ion intensities for each protein. Mass fraction for each protein was calculated by dividing the summed ion intensity for a given protein by the total ion intensity from all measured proteins. Mass fractions for the different sectors of cell composition depicted in Figure 1G were calculated by summing the mass fractions of all proteins associated with each sector. Mass fractions for the ethanol condition were taken from a previously published study (Lanz et al., 2024).

### Total proteome mass variance analysis

To define gross compositional change between two proteome measurements, the following equation was used:

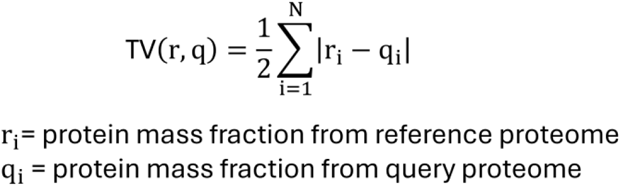

Total variation (TV) represents the summed change in protein mass fractions between two measured proteomes. The reference “proteome” in Figure 1E are the average mass fraction measurements of the three 0h WT samples. The TV measured between the three 0h WT samples was subtracted from the y-axis in Figure 1F so that the average 0h TV is set to 0.

### Principal component analysis

PCA analysis was performed in Python using the sklearn package. A dataframe was created that contained individual proteins as rows with columns corresponding to the relative protein concentration in each TMT channel (obtained from the median of all peptide measurements for a given protein). For the PCA analysis depicted in Figure S4, multiple independent experiments were combined by normalizing all measurements to the “0h” condition (log-phase growth in SCD media).

## Acknowledgements

We thank members of the Skotheim and Gottschling labs for helpful comments on the manuscript, and members of the Chan Zuckerberg Biohub’s Mass Spectrometry Platform for assistance in maintaining mass spectrometer performance. All proteomics data were acquired using Chan Zuckerberg Biohub’s Mass Spectrometry Platform.

## Supplementary Figures

**Figure S1.**
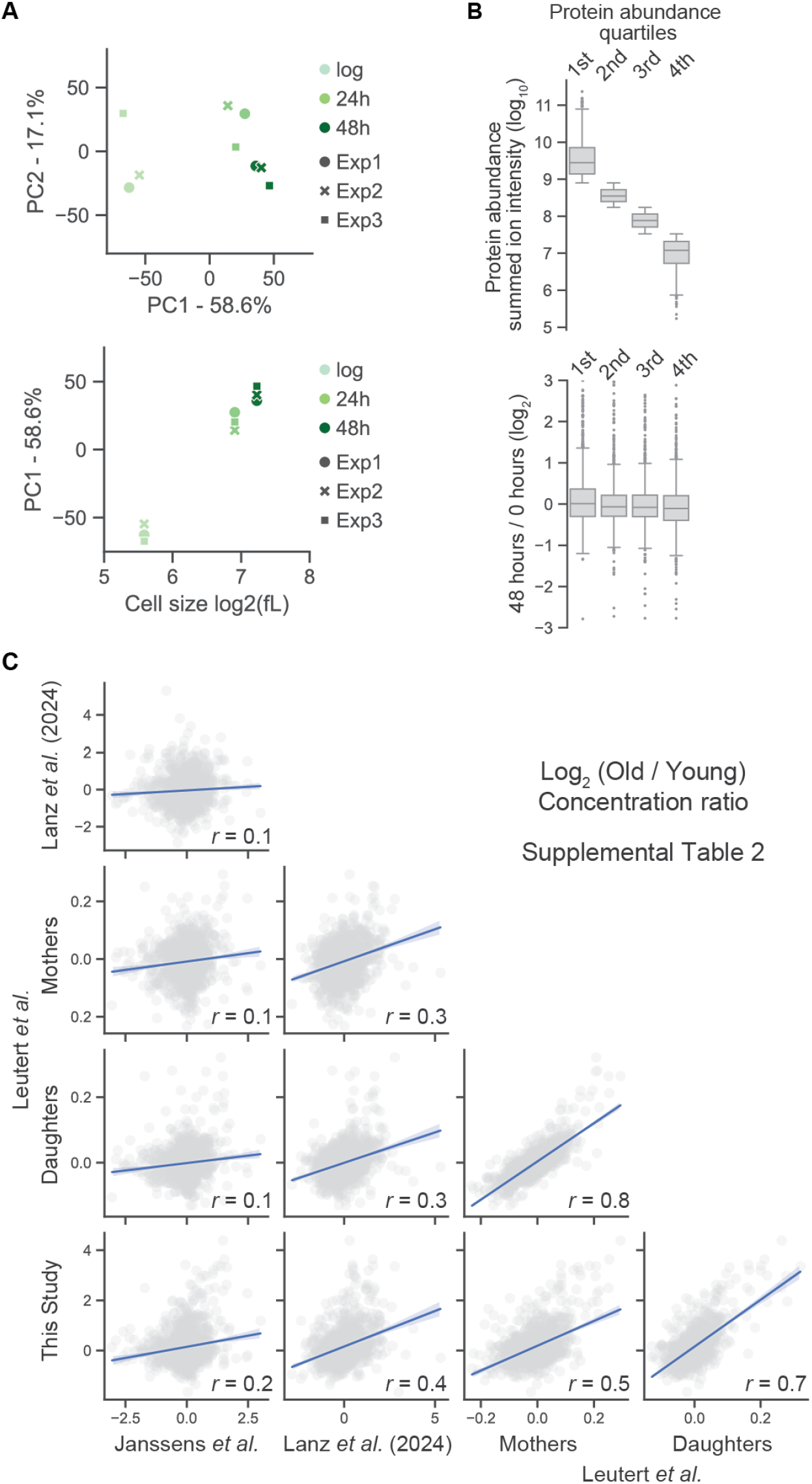
Analysis of principle components in the aged proteome and comparison with other studies. A) Principal component comparison of the three time points measured from three replicate time courses. Top panel: PC1 captures the majority of proteome variance and therefore represents a one-dimensional approximation of gross compositional change in the proteome. Bottom panel: This approximation mostly corresponds to the measured difference in cell size between the young and aged populations of yeast. B) To determine if protein abundance predicts age-associated behavior, the individual proteins were grouped into abundance quartiles based on their summed MS1-level peptide intensity (depicted in the upper plot). The bottom plot depicts the distribution of log_2_ concentration ratios (48 hours / 0 hours, n=3) for the proteins in each abundance quartile. Box region represents the median and interquartile range (IQR). Tails extend to 1.5x the IQR. C) Correlation grid of aging-associated proteome changes from four different published or pre-printed studies. See Table S2 for the “Old / Young” ratio metric used by each individual study. *r* denotes the pearson correlation coefficient. Blue line represents the regression with a 95% confidence interval.

**Figure S2.**
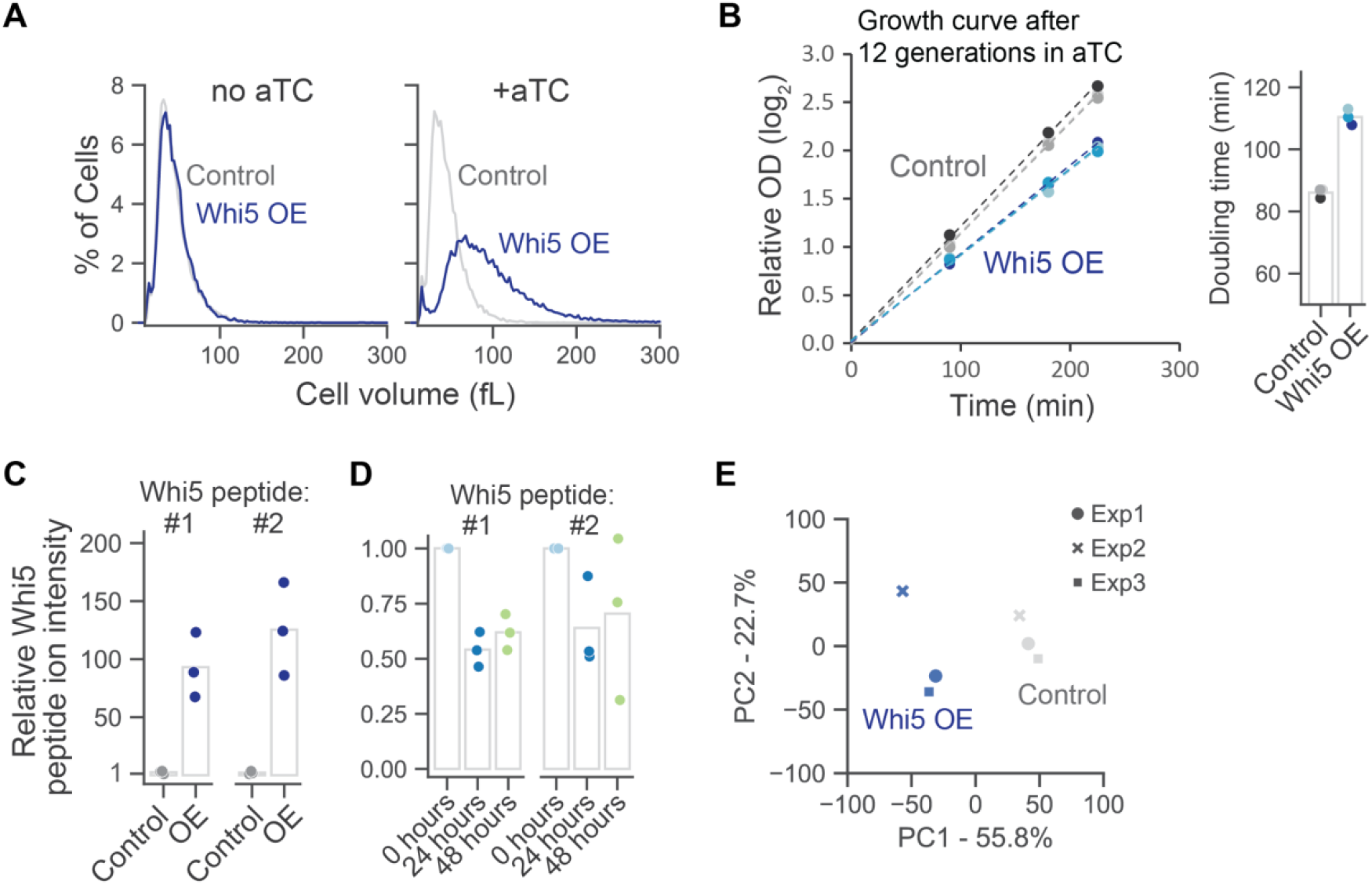
Cell size and growth in response to *WHI5* over-expression. A) Coulter counter measurements depicting the cell volume distributions for the experiment conditions depicted in Figure 2. Measurements were made twelve generations after adding anhydrotetracycline (aTc) to the culture flask to induce *WHI5* expression. B) Growth curve measuring the exponential growth rate of *WHI5* OE or control cell populations. Optical density (OD) is measured relative to the start of the time course (time = 0). The bar graph shows the calculated doubling time for three replicate cultures based on the depicted linear regressions. Different colors denote replicate cultures. C) Bar graph showing the TMT reporter ion intensities for the two measured Whi5 peptides. Each dot corresponds to a replicate *WHI5* OE experiment. For each experiment, Intensity is quantified relative to the control condition. D) Bar graph showing the TMT reporter ion intensities for the two measured Whi5 peptides. Each dot corresponds to a replicate MAD aging experiment. For each experiment, Intensity is quantified relative to the 0h condition. E) Principal component comparison of the three replicate *WHI5* OE experiments.

**Figure S3.**
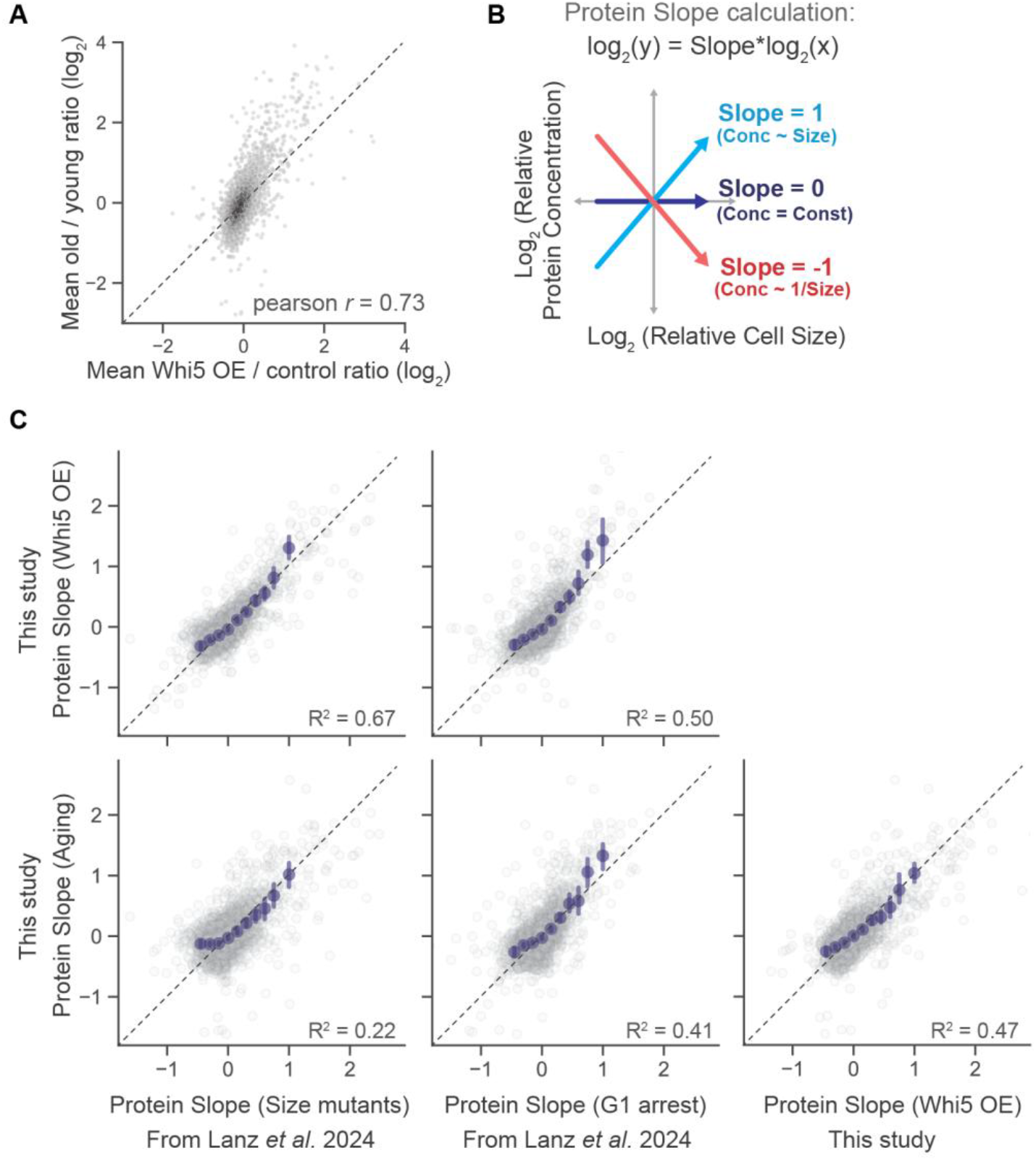
Comparison of protein concentration changes due to cell size increases and aging in diverse datasets. A) Correlation of (Old / young) and (*WHI5* OE / control) log_2_ ratios. Dotted line represents the identity line. Data density is shaded. B) Same schematic depicted in Figure 2. Derivation of the protein slope value. Protein slopes describe the size-scaling behavior of each individual protein. Proteins with a slope of 0 maintain a constant cellular concentration regardless of cell volume. A slope value of 1 corresponds to an increase in concentration that is proportional to the increase in volume and of −1 to dilution so it is proportional to 1/volume. C) Correlation of protein slope values derived using orthogonal experimental systems to change cell size (see Lanz et al., 2024 for more information on the “Size mutant” and “G1 arrest” experiments).

**Figure S4.**
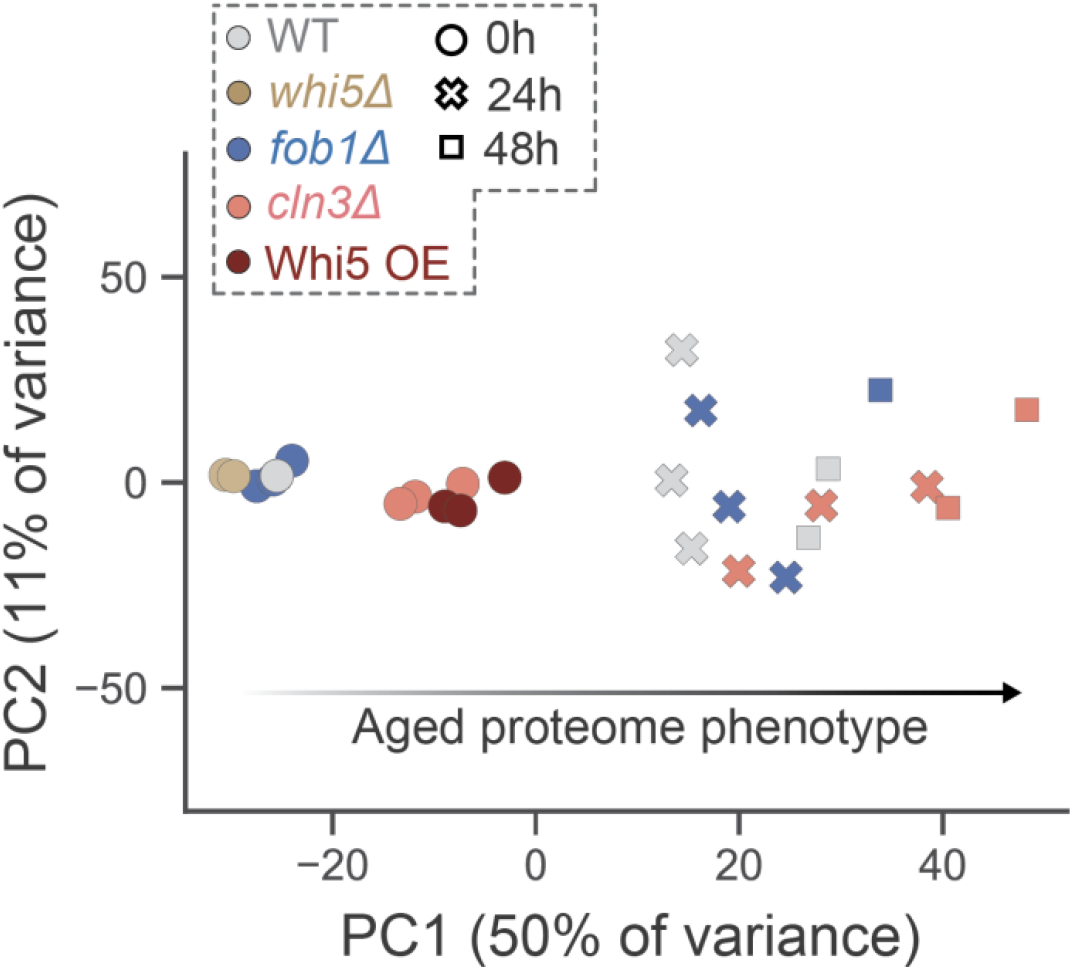
Principal component analysis showing how proteome composition changes with age in different cell size and longevity mutants. Principal component comparison of the aged proteome phenotypes of the indicated genotypes. Markers represent the time spent in the MAD. Three replicate MAD experiments with cell size and longevity mutants were normalized to 0h WT and plotted together. PC1 captures the majority of proteome variance and therefore represents a one-dimensional approximation of the progression of aging. *whi5Δ* proteomes were taken from a previously published binary comparison with WT (Lanz et al., 2024).

